# Differential Inhibition of Intra- and Inter-molecular Protease Cleavages by Antiviral Compounds

**DOI:** 10.1101/2023.06.28.546904

**Authors:** Jennifer S. Doherty, Karla Kirkegaard

## Abstract

Enteroviruses encode two protease active sites, in the 2A and 3C coding regions. While they target many host proteins, they first need to be excised from the viral polyprotein in which they are embedded. Polyprotein cleavage can occur either intra-molecularly (in *cis*) or inter-molecularly (in *trans*). Previous work suggested that antivirals targeting intra-molecular cleavages could generate inhibitory precursors that suppress the outgrowth of drug-resistant variants. Therefore, we wanted to evaluate enteroviral cleavage patterns to identify such obligate intra-molecular cleavages for drug target selection. Using translation extracts, we show that *cis* cleavage of the 2A protease N-terminal junction is conserved across three enteroviruses, while the mechanism for the N-terminal junction of 3C varies, with EV-D68 3C cleavage occurring in *cis* and poliovirus 3C cleavage occurring in *trans*.

Antiviral agents targeting proteases are often identified via their ability to block the cleavage of artificial peptide substrates. Here, we show that antivirals identified for their abilities to block inter-molecular cleavage can sometimes block intra-molecular cleavage of the protease from its polyprotein as well, but with widely varying efficacy. Additionally, we demonstrate that, for three enteroviral species, genomes defective in 2A protease activity suppress the growth of wild-type virus in mixed populations, supporting the hypothesis that preventing intra-molecular cleavage at the VP1·2A junction can create dominantly inhibitory precursors. These data argue that, to reduce the likelihood of drug resistance, protease-targeted antivirals should be evaluated for their ability to block intra-molecular polyprotein cleavages in addition to inter-molecular cleavage of other substrates.

**IMPORTANCE:** Most protease-targeted antiviral development evaluates the ability of small molecules to inhibit cleavage of model substrates. However, before they can cleave any other substrates, viral proteases need to cleave themselves from the viral polyprotein in which they have been translated. This can occur either intra- or inter-molecularly. Here, we show that, for poliovirus, Enterovirus D68 and Enterovirus A71, many of these cleavages are required to occur intra-molecularly. Further, we show that antivirals identified for their ability to block cleavage of artificial substrates can also block intra-molecular self-cleavage, but that their efficacy in doing so varies widely. We argue that evaluating candidate antivirals for their ability to block these cleavages is vital to drug development, because the buildup of uncleaved precursors can be inhibitory to the virus and potentially suppress the selection of drug-resistant variants.

## INTRODUCTION

Enteroviruses (EV) constitute a genus of positive-strand RNA viruses that includes significant human pathogens such as poliovirus, Rhinovirus, EV-A71, and EV-D68. They form part of the larger *Picornaviridae* family. Though poliovirus has been largely eradicated, vaccine-derived cases still emerge (1–5), and have recently caused outbreaks in Israel, the UK, and the US (6). EV-A71 can lead to severe neurological outcomes in children (7–9) and EV-D68 has been associated with acute flaccid myelitis among children in recent years (10–13). There are currently no vaccines or antivirals available for the treatment of these emerging enteroviruses.

All members of the enterovirus family share the same genomic architecture. These genomes are translated as polyproteins which must be processed into individual viral gene products by two virally encoded proteolytic activities. 3C and 3CD make most of the cleavages while 2A cleaves at the border between capsid protein VP1 and its own coding region (Fig. 1A). Both proteases also cleave many cellular targets to make the cell more hospitable for viral replication (14–17), including eukaryotic initiation factor 4G(eIF4G) by 2A, whose cleavage stimulates the preferential translation of viral proteins during infection (18). Though enteroviral proteases have been extensively studied as drug targets, no enteroviral protease inhibitors have yet been approved for clinical use. A deeper understanding of enterovirus biology could serve as a basis for more rational drug design for all viruses that utilize a polyprotein strategy.

**Figure 1.**
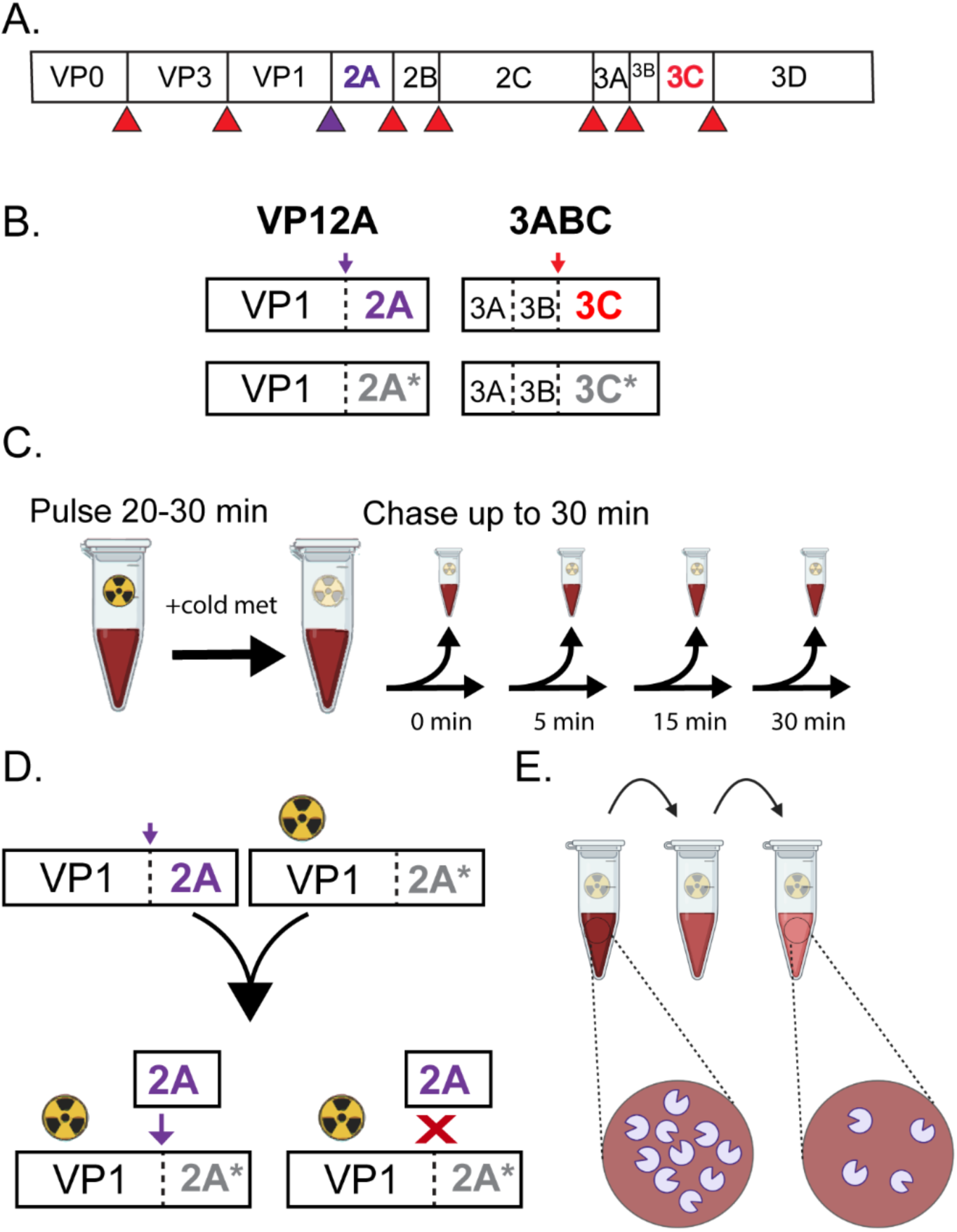
Design of protein processing assays. A) Enterovirus polyprotein showing cleavages made by 3C protease activity (red) and 2A protease activity (purple). B) Constructs used to program translation extracts from rabbit reticulocytes. 2A* and 3C* represent catalytically inactive mutant proteins 2A C109R and 3C C147R. C) Pulse chase experimental design D) *Trans* cleavage assay design. Catalytically active, unlabeled protease is combined with catalytically inactive protein that has been radioactively labeled. If *trans* cleavage can occur, the inactive precursor will be processed (left), and if it cannot no processing will occur (right) E) Dilution sensitivity assay design. If processing occurs in *cis*, the rate of the reaction should not change upon dilution.

Release of polyprotein-embedded proteases can occur either intra-molecularly, with a single molecule cleaving itself, or inter-molecularly, with one molecule cleaving a second molecule. These mechanisms can also be referred to as *cis* (intra-molecular) and *trans* (inter-molecular). In positive-stranded RNA virus groups such as flaviviruses (19, 20) and alphaviruses (21, 22), some of the polyprotein cleavages have been shown to occur exclusively in *cis*, enabling rapid excision from the polyprotein and providing a unique opportunity for antiviral development. RNA virus populations exist as a quasispecies (23). Therefore, drug-susceptible and newly minted drug-resistant variants coexist in the same cell and can interact. In dengue virus, mutant genomes encoding proteins that fail to make an obligate intra-molecular cleavage were shown to have an inhibitory effect on wild-type growth in mixed infections (20), arguing that blocking such cleavages could lead to the production of precursors that act as dominant inhibitors. Therefore, identification of intra-molecular cleavages in viral proteins will reveal promising substrates for antiviral targeting.

In the Enterovirus genus, poliovirus polyprotein processing has been the most intensively studied. While some work on poliovirus 2A has suggested its excision from the polyprotein occurs exclusively intra-molecularly (23, 24) other data suggested this proteolytic event could occur in *trans* (25, 26). For 3C and 3CD proteases, inter-molecular processing of polyprotein precursors has been extensively investigated (27–30), but the mechanism of processing for the 3C N-terminal junction itself is unresolved. Some work in poliovirus suggested that this processing can occur inter-molecularly (29, 31, 32), while work in ECMV, a non-enterovirus member of the *Picornaviridae*, showed that 3C processing was dilution insensitive, and therefore likely occurs intra-molecularly (33–35).

Protease inhibitors are often identified by screening for the ability to block the cleavage of small peptide substrates in *trans*. However, intra-molecular cleavages display different kinetics and may therefore serve as unique drug targets. In this work, we have identified several intra-molecular cleavages in the poliovirus and EV-D68 protein processing cascades and evaluated the efficacy of existing small molecules on both intra- and inter-molecular protease cleavages. We also show, for three different enteroviruses, that specifically blocking VP1-2A cleavage in a mixed infection can suppress the growth of all viruses in the same cell, even those that encode functional 2A protease. Thus, drugs inhibiting these intra-molecular cleavages may be especially effective due to their ability to suppress the emergence of resistant variants.

## RESULTS

To identify intra- and inter- molecular cleavages in enterovirus polyprotein processing, we developed constructs encoding short regions of the viral polyprotein that include the protease of interest. These constructs were used to program translation extracts (Fig. 1B) and the translation products were then used as substrates in pulse-chase reactions (Fig. 1C). To assay *trans* cleavage directly, constructs that encode catalytically inactive protease were translated in the presence of radioactive label. In the absence of enzymatic activity, the product is predicted to remain a full-length precursor (Fig. 1D, right). If *trans* processing can occur, the radioactively labeled precursor will be processed upon addition of an active, unlabeled protease (Fig. 1D, left). Thus, if no processing is observed, we can conclude that the precursor is cleaved exclusively intra-molecularly under these conditions. It is not possible to measure *cis* cleavage alone because multiple copies of a protein are translated, so *trans* cleavage is always a possibility. Therefore, to confirm the mechanism of *cis* cleavages, we can test the dilution sensitivity of the processing of wild-type construct. To this end, catalytically active construct translated with radioactive label is diluted during the chase period (Fig. 1E). Upon dilution, the rate of intra-molecular reactions should be unchanged while inter-molecular reactions are predicted to occur more slowly.

### Poliovirus 2A protease cleaves its N-terminal junction primarily intra-molecularly

To determine whether the excision of the N-terminus of 2A protease occurs in *cis* or in *trans*, we used a construct encoding only VP1·2A to program translation extracts. Representative images for the *trans* cleavage assay are shown in Figure 2A. Protein molecular weights are listed in Table S1. As expected, VP1·2A with a mutation in the 2A active site was not processed (left panel). In the presence of catalytically active, unlabeled VP1·2A, processing was incomplete (center panel). In contrast, self-processing of the wild-type construct, which can occur either in *cis* or in *trans*, was rapid and complete (right panel). These data are quantified in Figure 2B and argue that the primary mode of cleavage for the VP1·2A junction is intra-molecular under these conditions. This is confirmed by the results of the dilution-sensitivity assay, which shows that the rate of precursor processing does not change upon dilution (Fig. 2 C, D). To verify that the 2A protease can cleave in *trans* under the assay conditions, we immunoblotted for eIF4G, which is naturally present in the translation extracts. In poliovirus-infected cell lysates, eIF4G cleavage was complete by five hours post infection (Fig. S1, lane 5). In the programmed translation reactions, only the catalytically active 2A construct could cleave eIF4G (Fig. S1, lane 2).

**Figure 2.**
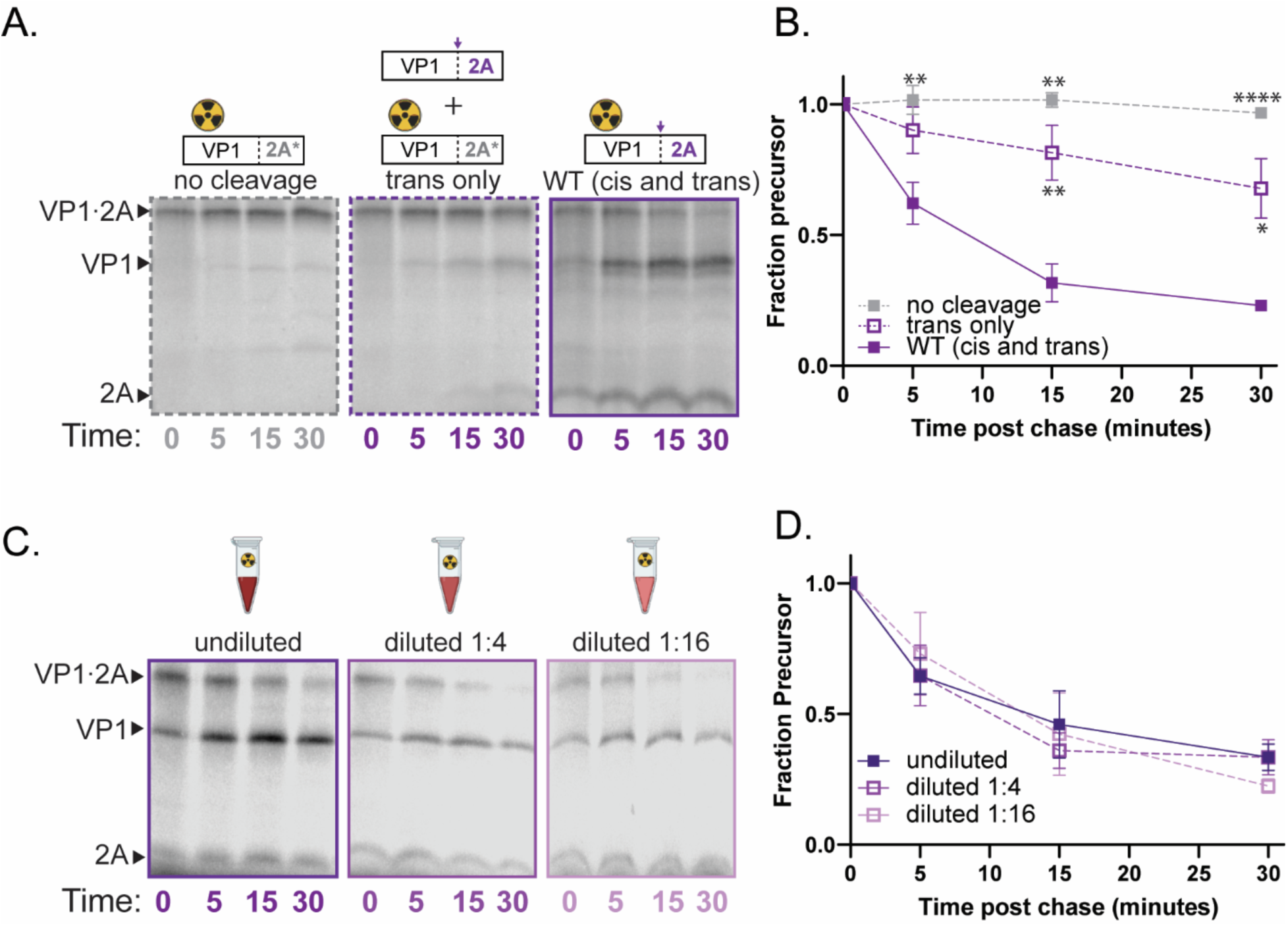
Poliovirus 2A processing occurs in *cis*. A) Representative gels for poliovirus 2A *trans* cleavage assay. 2A* represents the C109R active site mutation. No cleavage (left): C109R construct with unlabeled WT 3ABC (negative control). *trans* only (middle): C109R construct with unlabeled WT VP1·2A. WT (*cis* and *trans*)(right): catalytically active VP1·2A. B) quantification of *trans* cleavage assay. Fraction precursor calculated by dividing the intensity of the VP1·2A band by the sum of the VP1·2A and the VP1 band intensities and is normalized to the value at time 0. C) representative gels for poliovirus 2A dilution sensitivity assay D) quantification of VP1·2A dilution sensitivity assay. Data quantified as in B.

### Poliovirus 3C protease cleaves its N-terminal junction primarily inter-molecularly

To determine whether the excision of the N-terminus of 3C protease occurs in *cis* or in *trans*, we also monitored its processing by pulse-chase reactions in translation exacts. Representative gels for the *trans*-cleavage assay are shown in Figure 3A and quantified in Figure 3B. Protein molecular weights are listed in Table S1. Processing of 3ABC occurred primarily between 3B and 3C. The rate of processing was identical in the *trans* only reaction and in the wild-type reaction in which both *cis* and *trans* cleavage could occur (Fig. 3 A,B). Additionally, the rate decreased with increasing dilution (Fig. 3C, D). Both findings are characteristic of an inter-molecular cleavage pattern. In all assays with labeled wild-type precursor (Fig. 3A, right; Fig 3C), a band corresponding to 3BC is also produced. This band is not seen in the *trans* cleavage condition (Fig 3A, middle). Thus, cleavage from 3ABC to 3BC is made in these extracts intra-molecularly, but only inefficiently. An additional slightly larger band was also observed on many of our poliovirus 3ABC gels (Fig. 3A, C; ǂ) and is likely the result of inappropriate translation initiation that sometimes occurs in rabbit reticulocyte lysates (36).

**Figure 3.**
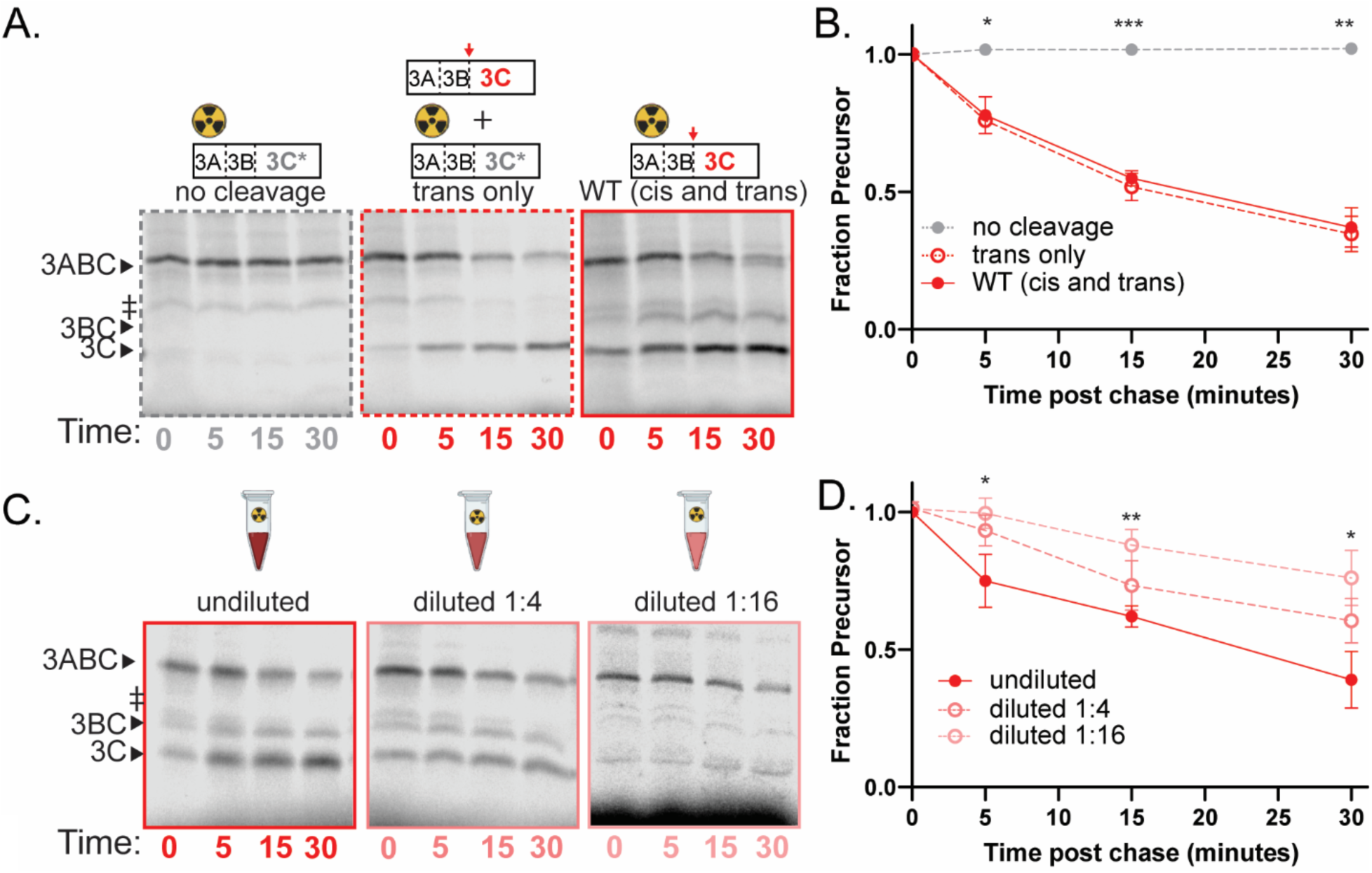
Poliovirus 3C processing occurs primarily in *trans*. A) Representative gels for poliovirus 3C *trans* cleavage assay. 3C* represents the C147R active site mutation. No cleavage (left): poliovirus 3C C147R with unlabeled WT VP1·2A (negative control); *trans* only (middle): 3C C147R with unlabeled catalytically active 3ABC. WT (*cis* and *trans*) (right): catalytically active 3ABC. ǂ represents an aberrant translation product due to internal translation initiation.(36) B) quantification of *trans* cleavage assay. Fraction precursor calculated by dividing the intensity of the 3ABC band by the sum of the 3ABC and the 3C band intensities and is normalized to the value at time 0. C) representative gels for poliovirus 3C dilution sensitivity assay D) quantification of dilution sensitivity assay. Data quantified as in B.

### EV-D68 2A and 3C processing occur primarily intra-molecularly

To determine how conserved these mechanisms are across the enterovirus genus, we established the *cis* and *trans* cleavage efficiencies for EV-D68 2A and 3C proteases. In the *trans-*cleavage assay for 2A protease, no inter-molecular processing of EV-D68 VP1·2A was observed although processing occurred rapidly for the active precursor (Fig. 4A,B). Exclusively intra-molecular cleavage was also observed for the VP1·2A precursor of Enterovirus A71 (Fig. S2), arguing that rapid intra-molecular cleavage between the capsid proteins and 2A protease is likely to be conserved across the Enterovirus genus.

**Figure 4.**
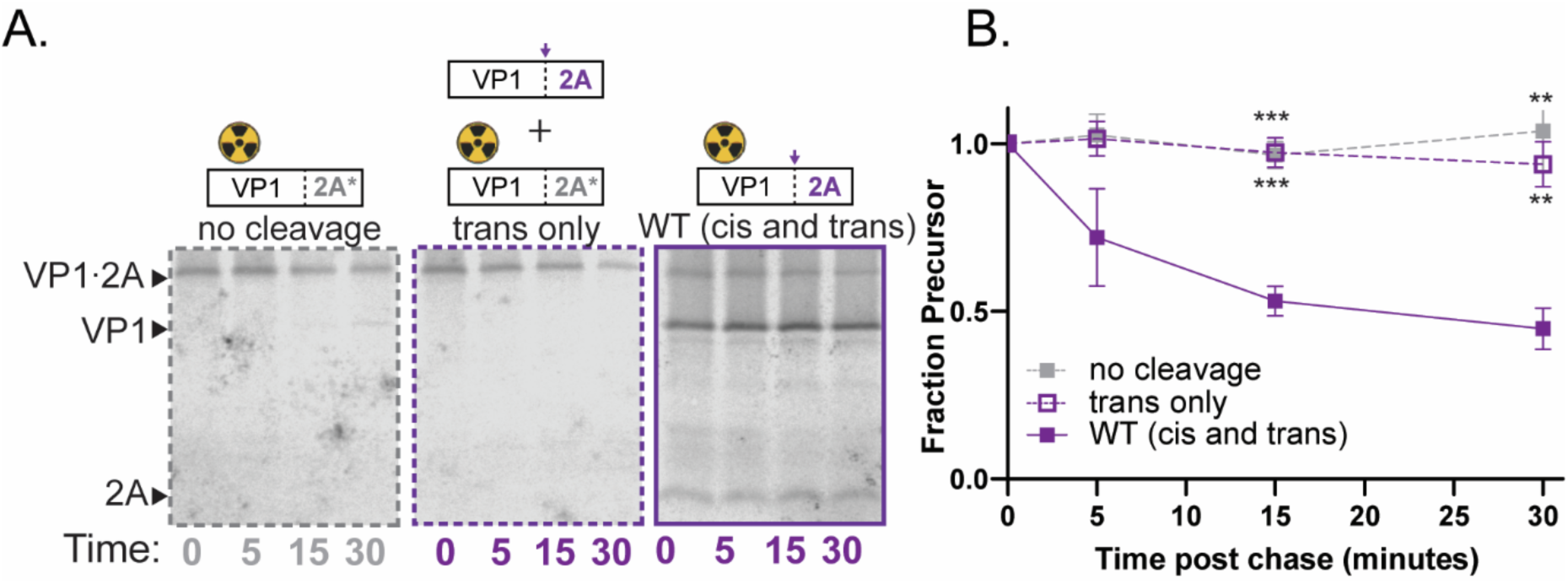
EV-D68 self-processing by 2A occurs in *cis*. Representative gels A) and quantification B) for EV-D68 VP1·2A *trans* cleavage assay. 2A* represents the C107R active site mutation. No cleavage: EV-D68 2A C107R with unlabeled 3ABC (negative control). *Trans* only: EV-D68 2A C107R with unlabeled catalytically active VP1·2A; WT (*cis* and *trans*): EV-D68 catalytically active VP1·2A. Data quantified as in Figure 2.

The processing of 3ABC precursors, however, differed between EV-D68 and poliovirus. Unlike the effective *trans* cleavage observed with poliovirus 3ABC (Fig. 3A, B), EV-D68 3ABC processing in the *trans*-only cleavage condition occurred much more slowly and incompletely (Fig. 5A,B), implying that the cleavage occurs primarily intra-molecularly. The dilution-sensitivity assay, which showed that the rate of 3ABC processing did not change upon dilution (Fig. 5C, D), corroborates this interpretation. Because 3C and 3CD exhibit different substrate recognition properties, we wanted to determine whether 3ABCD precursor processing exhibited different kinetics from 3ABC processing. Extracts programmed with 3ABCD constructs also showed a strong preference for *cis* processing, with wild-type cleavage so rapid that little full-length precursor was observed at the beginning of the chase period (Fig. 5E,F).

**Figure 5.**
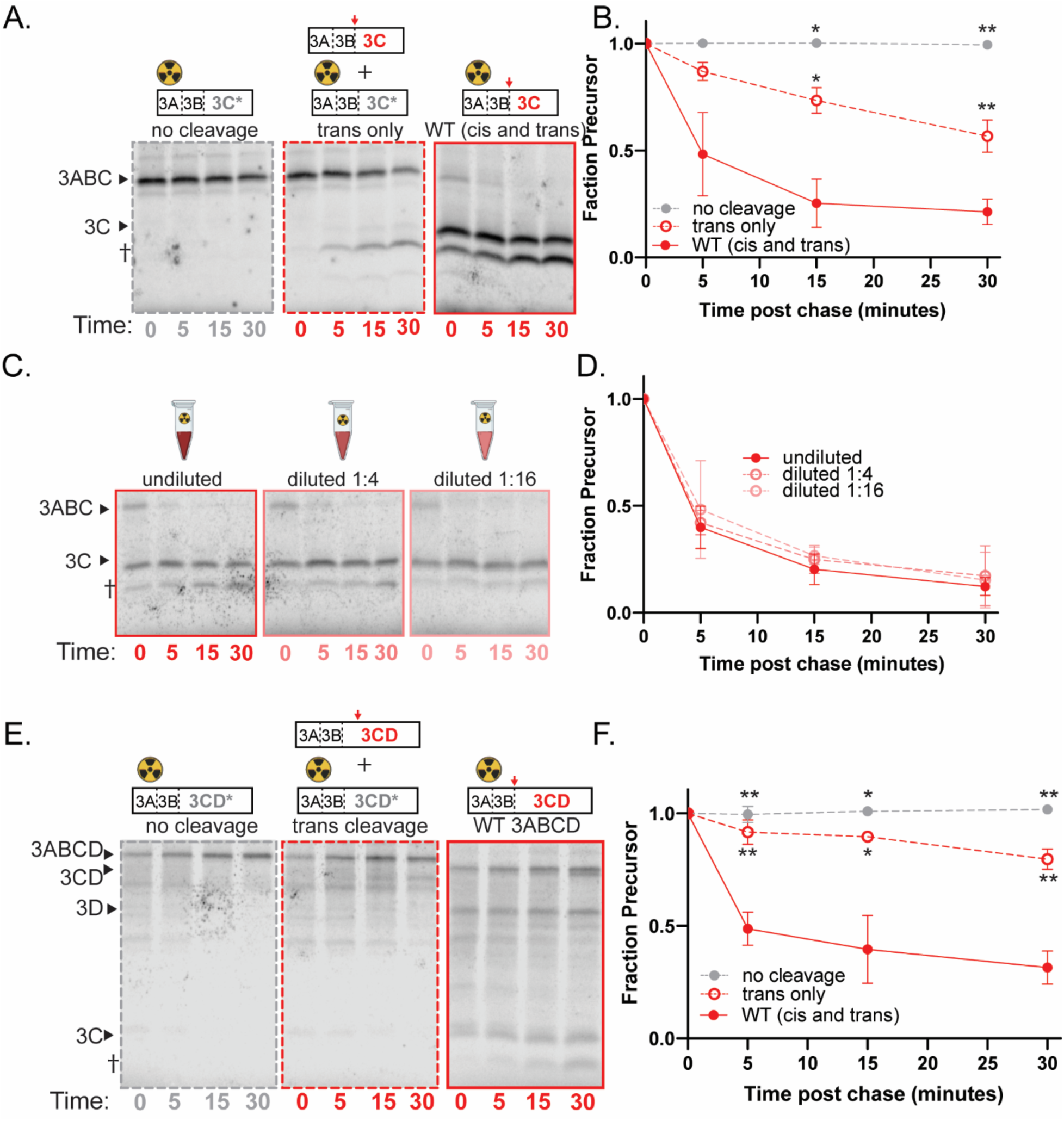
EV-D68 3C protease self-processing occurs in cis. Representative gels (A) and quantification (B) of EV-D68 3ABC *trans* cleavage assay. 3C* represents the C147R active site mutation. No cleavage: EV-D68 3C C147R with unlabeled VP1·2A (negative control). *Trans* cleavage: EV-D68 3C C147R with unlabeled catalytically active 3ABC; WT (cis and trans): EV-D68 catalytically active 3ABC. † indicates unidentified cleavage product. Fraction precursor calculated by dividing the intensity of the 3ABC band by the sum of the 3ABC, 3C, and † band intensities and is normalized to the value at time 0. C) representative gels of EV-D68 3ABC dilution sensitivity assay. D) quantification of EV-D68 3ABC dilution sensitivity assay. Quantified as in B. E) Representative gels of 3ABCD *trans* cleavage assay. 3CD* represents the 3C C147R catalytically inactive mutant. No cleavage: EV-D68 3C C147R construct with unlabeled VP1·2A (negative control); *trans* only: EV-D68 3C C147R with unlabeled catalytically active 3ABCD. WT (cis and trans): EV-D68 catalytically active 3ABCD. F) Quantification of EV-D68 3ABCD *trans* cleavage assay. Fraction precursor calculated by dividing the intensity of the 3ABCD band by the sum of the 3ABCD, 3CD, and 3D band intensities and is normalized to the value at time 0.

EV-D68 3ABC and 3ABCD processing produced an additional band of approximately 16 kDa that does not correspond to the size of any identified 3ABC cleavage products and is of unknown origin (Fig. 5A,C, Fig. S3). The production of this band requires 3C protease activity, as is it not present in translation extracts programmed with catalytically inactive 3C (Fig. 5A,C) or when 3C is inhibited by an antiviral drug (Fig. 8A). Interestingly, this unknown band is the major cleavage product when EV-D68 3ABC cleavage is forced to occur in *trans*, highlighting the importance of intra-molecular cleavage reactions for proper polyprotein processing.

### Antivirals have varying efficacy in inhibiting intra-molecular cleavages

Proteases of positive-strand viruses have two types of substrates during infection: host proteins, which must be cleaved inter-molecularly, and viral polyproteins, which can be cleaved either intra-molecularly or inter-molecularly. Few studies have been done to correlate protease inhibitor efficacy against virus with the type of cleavage that is being inhibited. To determine how effectively the different cleavage events that occur during poliovirus and EV-D68 infections are inhibited by existing protease inhibitors, we investigated the effects of 2A inhibitors telaprevir and zVAM.fmk and 3C inhibitor rupintrivir on the inter- and intra-molecular cleavages described here.

Telaprevir, originally identified as an inhibitor of hepatitis C virus NS3/NS4A (37), was recently shown to be a potent and specific inhibitor of EV-D68 2A protease, with reported EC_50_ values for EV-D68 viral growth ranging from 0.5-2.0 µM (38). To compare the ability of this small molecule to inhibit cleavage of inter-molecular targets, for which it was identified, with its ability to inhibit intra-molecular self-processing, an EV-D68 VP1·2A construct was translated in the presence of increasing concentrations of telaprevir (Fig. 6A, top). To determine the effect of telaprevir on inter-molecular cleavage of host eIF4G, a construct encoding 2A protease alone was translated without radioactive methionine, and the proportion of intact eIF4G within the translation extract was ascertained via immunoblot (Fig. 6A, bottom). At 1 µM telaprevir, the maximum concentration we could test in our translation extracts, the intra-molecular processing of VP1·2A was only 20% inhibited, while the inter-molecular processing of eIF4G was fully inhibited (Fig. 6B). Based on these data, the approximate IC_50_ for the intra-molecular cleavage of VP1·2A is around 4 µM, while inter-molecular cleavage of eIF4G is more potent, with an IC_50_ of approximately 200 nM (Table 1). To characterize the inhibition of EV-D68 VP1·2A self-processing by telaprevir further, pulse-chase experiments were performed to monitor the kinetics of precursor processing in the presence and absence of 1µM of the drug (Fig. 6C). In the presence of telaprevir, the rate of VP1-2A self-processing was significantly decreased for the wild-type construct (Fig. 6D), but not for a telaprevir-resistant mutant (Fig. S4), confirming that the reduction in VP1·2A self-processing, although modest, is due to inhibition of 2A activity.

**Figure 6.**
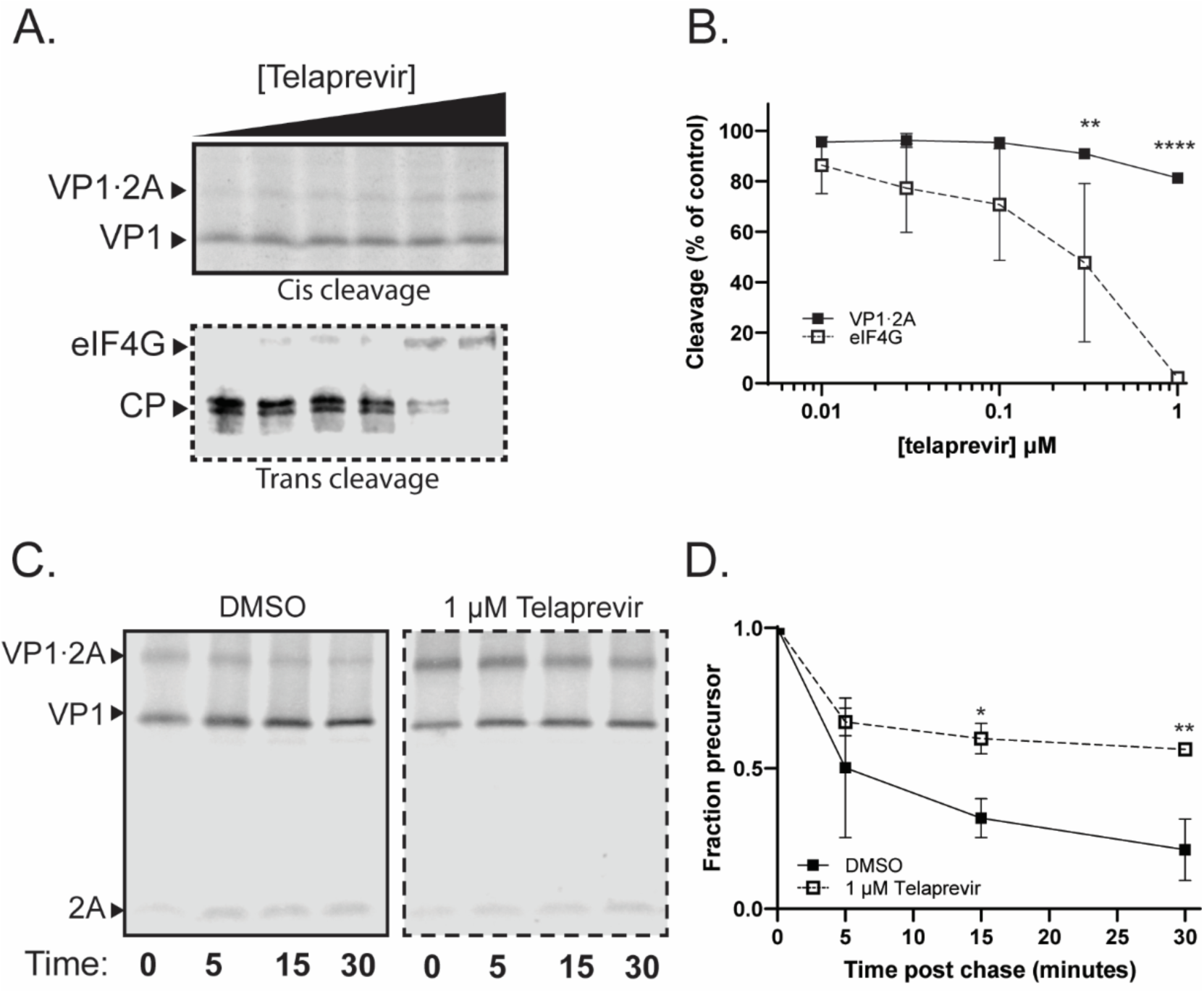
Telaprevir can inhibit the intramolecular cleavage of EV-D68 2A. A) Representative gels of self-processing (top panel) and eIF4G cleavage (bottom panel) by EV-D68 2A in the presence of increasing concentrations of Telaprevir. Telaprevir was added at the initiation of translation. CP, eIF4G cleavage products B) quantification of telaprevir concentration curves. Percent cleavage normalized to the amount of cleavage in the DMSO condition (first lane) (C and D) Representative gels (C) and quantification (D) of EV-D68 VP1·2A pulse chase experiment in the presence of telaprevir. Data quantified as in Figure 2.

**Table 1.** Approximate IC50 values for intra- and inter-molecular cleavages by protease targeted antivirals.

| Drug | Cleavage | IC<sub>50</sub><br>(μM) | 95% CI<br>(μM) |
| --- | --- | --- | --- |
| **Telaprevir** | **EV-D68 VP1·2A** (intra-molecular) | 3.96 | 3.17-5.12 |
|  | **eIF4G** (inter-molecular) | 0.19 | 0.099-0.356 |
| **zVAM.fmk** | **Poliovirus VP1·2A** (intra-molecular) | 1570 | 1320-1890 |
|  | **eIF4G** (inter-molecular) | 18.9 | 12.6-27.3 |
| **Rupintrivir** | **EV-D68 3ABC** (intra-molecular) | 0.14 | 0.146-0.193 |
|  | **Poliovirus 3ABC** (inter-molecular) | 0.76 | 0.584-0.988 |

Peptide inhibitor zVAM.fmk was initially identified as a rhinovirus protease inhibitor. It was shown to inhibit both rhinovirus VP1·2A precursor processing and eIF4G cleavage with optimal effects at 200µM (39). To determine whether zVAM.fmk also inhibits poliovirus 2A protease, translation extracts programmed with poliovirus VP1·2A were incubated with increasing concentrations of zVAM.fmk (Fig. 7A). Again, intra-molecular cleavage of VP1·2A, while significantly inhibited by zVAM.fmk, was much less affected than the inter-molecular cleavage of eIF4G (Fig. 7 A,B). The calculated IC_50_ for intra-molecular VP1·2A cleavage is approximately 1.6 mM, while inter-molecular eIF4G cleavage was much more effective, with an IC_50_ of approximately 20µM (Table 1). Pulse-chase experiments also revealed significant inhibition of VP1·2A *cis* cleavage at high zVAM-fmk concentrations (Fig. 7C,D) confirming the compound’s efficacy but low potency for this intra-molecular reaction.

**Figure 7.**
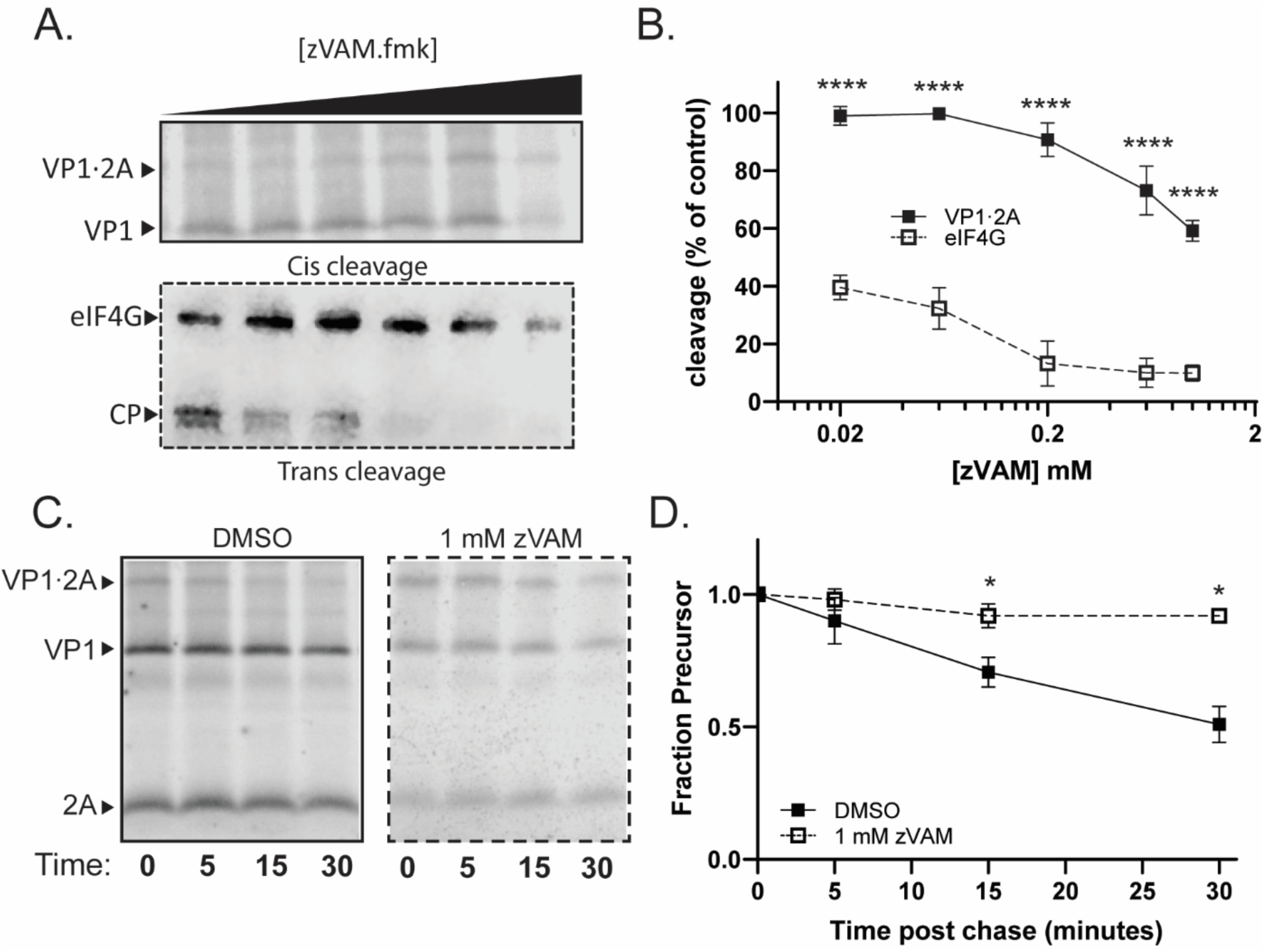
zVAM.fmk can inhibit self-processing of poliovirus 2A. A) Representative gels of self-processing (top panel) and eIF4G cleavage (bottom panel) by poliovirus 2A in the presence of increasing concentrations of zVAM.fmk. zVAM.fmk was added at the initiation of translation. CP, eIF4G cleavage products. B) quantification of zVAM.fmk concentration curves. Percent cleavage normalized to the amount of cleavage in the DMSO condition (first lane). (C and D) Representative gels (C) and quantification (D) of poliovirus VP1·2A pulse chase in the presence of 1 mM zVAM.fmk

Rupintrivir was identified as an inhibitor of rhinovirus 3C protease (40), but has since been shown to inhibit the enzymatic activity of a wide variety of viral 3C and 3C-like proteases (41–45). It is an efficacious inhibitor of viral growth as well: the reported EC_50_ for EV-D68 isolates ranges from 2-3 nM (42), while for poliovirus the reported EC_50_ ranges from 5-40 nM (46). In translation extracts, we observed near complete inhibition of both EV-D68 and poliovirus 3ABC processing, albeit at different concentrations (Fig. 8 A-D). The IC_50_ for intra-molecular cleavage of EV-D68 3ABC is approximately 140 nM, while inhibition of the inter-molecular cleavage of poliovirus 3ABC is approximiately 750 nM (Table 1). In this case, the intra-molecular cleavage of EV-D68 3ABC is approximately five-fold more susceptible to rupintrivir than the inter-molecular cleavage of poliovirus 3ABC, which correlates with the relative effectiveness of rupintrivir against viral replication. Upon pulse-chase, we observed essentially 100% inhibition for both EV-D68 intra-molecular processing (Fig. 8 E,F) and poliovirus inter-molecular processing (Fig. 8 G,H).

**Figure 8.**
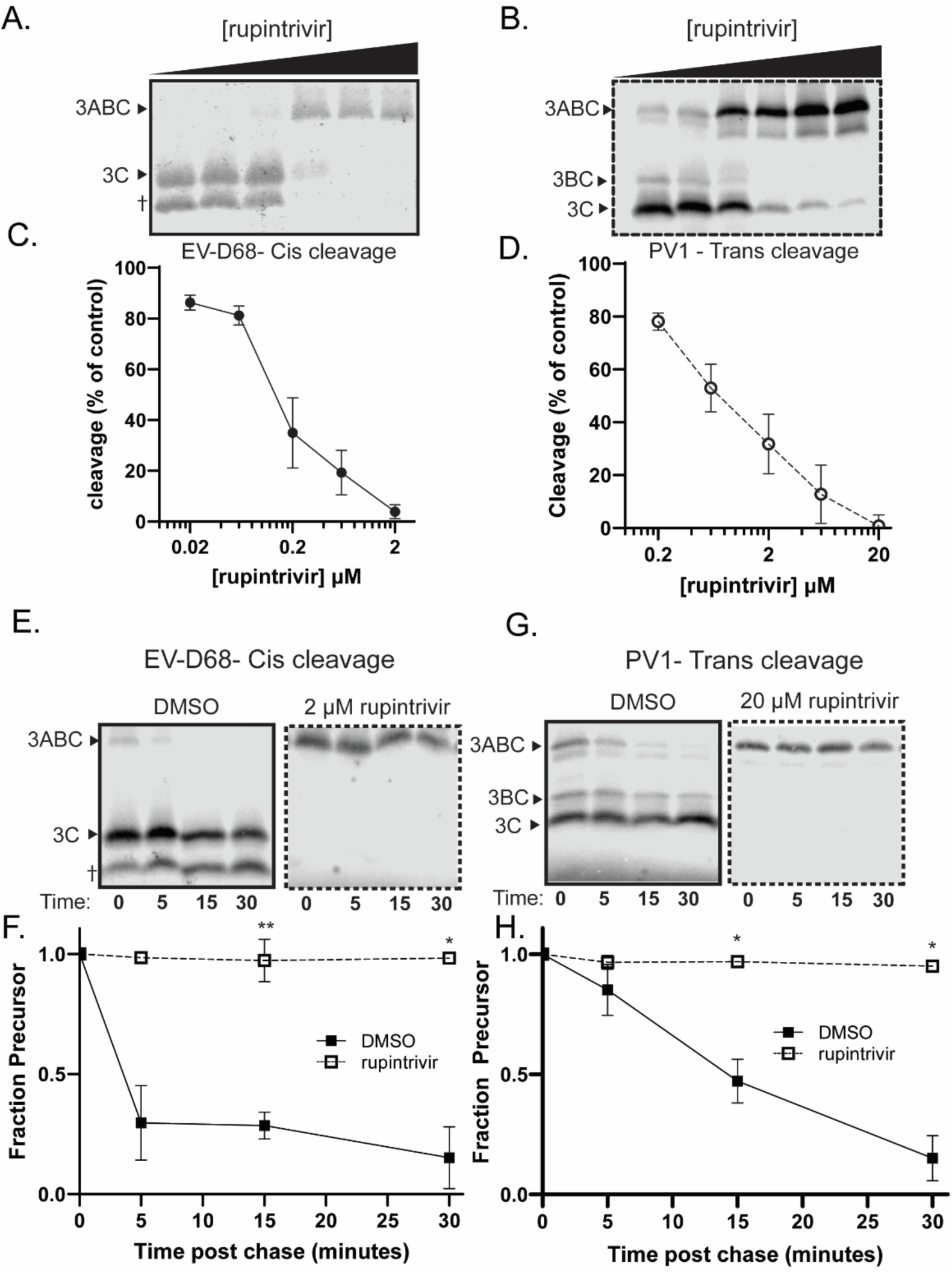
Rupintrivir inhibits 3ABC self-processing in poliovirus and EV-D68. (A and B) representative gels of rupintrivir concentration curve in EV-D68 (A) and Poliovirus (B). Rupintrivir was added at the initiation of translation. (C and D) quantification of A and B. Percent cleavage normalized to the amount of cleavage in the DMSO condition (first lane) (E and F) Representative gel and quantification of EV-D68 pulse chase experiment in the presence of 2 µM rupintrivir. (G and H) Representative gel and quantification of poliovirus pulse chase experiment in the presence of 20 µM rupintrivir.

### Inhibition of 2A protease is genetically dominant

For most antivirals, the target simply loses function in the presence of the drug and resistant variants will outcompete drug-susceptible variants. However, antivirals that specifically block intra-molecular cleavages could suppress the emergence of drug-resistant variants, if blocking such cleavages led to the accumulation of unprocessed precursors that interfered with the growth of more fit viruses (20, 47). We consider such cleavages to be potential dominant drug targets, because inhibited drug-susceptible proteins can prevent the amplification of drug resistant genomes, masking the resistance phenotype (Fig. 9A).

**Figure 9.**
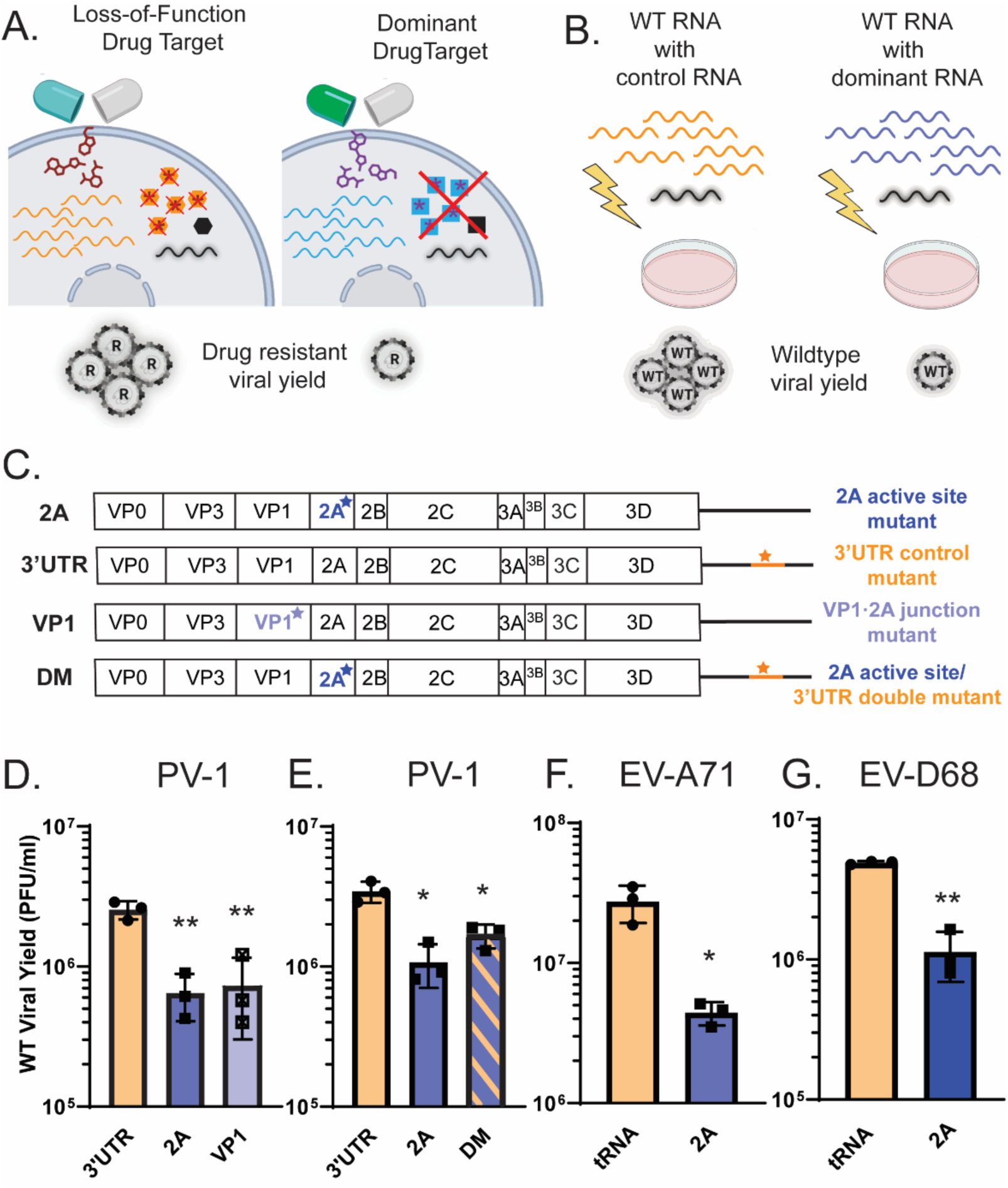
Genomes with inactive 2A are dominant over those with wild-type 2A. A) Model of dominant drug targets. In a loss of function drug target, viral products from drug susceptible genomes (orange hexagons) have no effect on a newly arisen drug resistant variant (black hexagon), so the drug resistant variant replicates freely and takes over the population. For a dominant drug target, the drug susceptible viral protein (blue square) has a toxic effect on drug resistant viral products (black square) and the replication of the drug resistant variant is suppressed. B) Cotransfection experimental design to model a scenario in which a drug resistant variant has arisen surrounded by drug susceptible variants. WT viral genomes (black) are cotransfected with an excess of either control RNA (orange) or potentially dominant mutant RNA (blue) via electroporation. Yield of WT viral RNA is quantified. C) Viral genomes used in cotransfection experiments. 3’UTR is replication incompetent and known to be non-dominant, and is therefore used as a control (D and E) Cotransfection of WT poliovirus with indicated mutant RNA. 3’UTR, 3’UTR ΔGUA3 mutant. 2A, 2A C109R mutant. VP1, VP1 Y302P mutant. DM, 2A C109R 3’UTR ΔGUA3 double mutant F) Contransfection of WT EV-A71 with 2A C110A mutant. tRNA used as a control for transfection of excess RNA. G) Contransfection of WT EV-D68 with 2A C107R mutant. tRNA used as a control for transfection of excess RNA. All titers show the amount of WT virus formed in a mixed transfection with the indicated mutant. Poliovirus transfections analyzed by one way ANOVA, compared to 3’UTR condition. EV-A71 and EV-D68 transfections analyzed by students t test.

To test for dominance, we can recreate the circumstances of an intracellular quasispecies by co-transfecting cells with wild-type viral RNA and an excess of mutant viral RNA (Fig. 9B). In this case the RNA with a mutation that inactivates the 2A protease is conceptually equivalent to an RNA genome whose 2A protein is rendered unfit by an antiviral. If infectious wild-type viral growth is inhibited by the products of these genomes, it suggests that 2A protease inhibition may lead to the generation of dominant inhibitors (Fig. 9B). Because RNA transfection efficiency is sensitive to the total amount of RNA, controls include the co-transfection of either a 10-fold excess of yeast tRNA or of a defective viral genome known not to have a negative effect on wild-type growth due to its mutation in the 3’ untranslated region.

In poliovirus, variants that fail to make the cleavage between VP1 and 2A, either because of a mutated protease active site or a mutated cleavage site, suppress the growth of wild-type viral genomes, while genomes with mutations in the 3’ untranslated region did not (Fig. 9 C,D; (48)). However, unlike the 3’UTR mutant genomes, 2A mutant genomes can undergo RNA replication (Fig. S5). To test whether the genetic dominance of the 2A mutant genomes was due to their ability to amplify their RNA, a defective genome containing both 2A and 3’ UTR mutations was subject to our co-transfection paradigm. The double-mutant genomes were also found to be dominant inhibitors of wild-type viral growth (Fig. 9E), arguing that this mechanism of inhibition is distinct from that of defective interfering RNAs, which require genome replication to be inhibitory (49).

These experiments demonstrate that the trans-dominance of poliovirus genomes encoding mutant 2A function correlates with lack of cleavage at the VP1·2A junction. This correlation is extended by the observation that both EV-A71 and EV-D68 genomes with 2A mutations suppress growth of the corresponding wild-type viruses (Fig. 9 F,G). Thus, we hypothesize that dominant inhibition of wild-type enterovirus genomes by 2A mutant genomes is due to the accumulation of a VP1·2A containing precursor whose processing cannot be accomplished in *trans*.

## DISCUSSION

Though viral proteases perform a variety of tasks during infection, their most immediate task is to cleave themselves from the polyprotein in which they are embedded. The timing and mechanism of these cleavages can play a regulatory role, controlling which viral protein products are present and at which proportions at various times during infection. Mechanistically, these cleavages can occur either intra-molecularly or inter-molecularly. Generally, intra-molecular cleavages occur more rapidly and completely than inter-molecular cleavages, and polyprotein cleavages that occur preferentially intra-molecularly could therefore be especially crucial for successful viral amplification.

In this study, we demonstrated that, for poliovirus, EV-A71 and EV-D68, the polyprotein cleavage between VP1 and 2A occurs almost exclusively intra-molecularly. This may represent a mechanism for the virus to quickly release the capsid proteins, which are large and difficult to fold (50) from the nascent polypeptide. In fact, mutations in 2A protease that slow the rate of polyprotein processing can partially rescue defects caused by malfunctioning capsid proteins (51), further highlighting the interplay between 2A function and capsid function. Many picornavirus groups that do not encode a 2A protease still have a mechanism to rapidly separate VP1 and 2A, for example by the addition of a ribosomal skipping mechanism in Aphthoviruses (52, 53), potentially pointing to a strong selection pressure to ensure this separation.

Unlike 2A cleavage, the mechanism of 3C cleavage differed across enteroviruses, with the 3B/3C cleavage occurring inter-molecularly in poliovirus, and exclusively intra-molecularly in EV-D68. This could represent an adaptation of EV-D68 to growth at lower temperatures, allowing the virus to maintain rapid self-cleavage kinetics at the 3B-3C junction. However, because only these two viruses were examined, it is difficult to draw more generalized conclusions about the biological function of these two different mechanisms. Given that the 3C protease primary sequence is actually slightly more conserved across the enterovirus genus than the 2A protease primary sequence(54), the fact that poliovirus and EV-D68 3C have different mechanisms of polyprotein excision is unexpected. Determining the mechanism of excision for other enteroviruses and the structures of protein precursors will help shed light on these observations.

In addition to a unique mechanism of polyprotein excision, EV-D68 3C is also unique in that we observed the production of an unidentified protein product of approximately 16 kDa resulting from self-processing of precursors containing 3C. This band appeared only when active cleavage could occur, demonstrating that it is a cleavage product rather than an aberrant translation product. We were unable to identify any secondary cleavage sites in the 3ABC precursor that could result in the production of this band. However, it is the major cleavage product when 3ABC processing is forced to occur in *trans*. Therefore, blocking the intra-molecular cleavage of EV-D68 3C could disrupt polyprotein processing both by causing a precursor to accumulate and by forcing that precursor to be processed incorrectly, providing multiple avenues through which toxic precursors could be produced. More work is needed to determine the identity of this protein and ascertain its relevance during infection.

We also examined the ability of antiviral drugs to block intra-molecular cleavages. Often, protease inhibitors are identified by their ability to inhibit the cleavage of small artificial peptides in *trans*. We wanted to determine how currently available antiviral compounds affect polyprotein processing as well, especially the intra-molecular cleavages. We saw that different inhibitors exhibited varying efficacy in blocking intra-molecular compared to inter-molecular cleavages. For example, poliovirus 2A inhibitor zVAM.fmk showed around 100-fold, and EV-D68 2A inhibitor telaprevir around five-fold, greater efficiency in blocking *trans* than *cis* cleavages. Rupintrivir was actually more efficient at blocking the *cis* cleavage of the 3AB.C junction of EV-D68 than the *trans* cleavage of the cognate junction of poliovirus 3ABC. These data imply that some, but not all, protease inhibitors identified for their ability to inhibit *cis* cleavages can also effectively inhibit *trans* cleavages. Therefore, evaluating candidate antivirals for their ability to inhibit intra-molecular cleavages as well as inter-molecular ones could help in prioritizing development efforts towards candidates that are more likely to be successful.

This becomes especially important considering our co-transfection findings. We showed that genomes encoding defective 2A protease are dominant over wild-type genomes in mixed infections. Due to the error-prone nature of the viral RNA-dependent RNA polymerase, many variants arise spontaneously during enterovirus infection and are subject to the crucible of selection. Though most will fail to propagate, some will encode useful features to the virus such as drug resistance. Our data argue that drugs blocking the intra-molecular cleavage between VP1 and 2A could potentially suppress the growth of resistant variants through the same mechanism that 2A mutant genomes suppress the growth of wild-type genomes in cotransfections. Therefore, drug development efforts targeted towards 2A should be especially fruitful because antivirals that target intra-molecular cleavage by 2A could help to evade the widespread problem of drug resistance.

## ACKNOWLEDGEMENTS

We thank Bert Semler, Tim Skern and Gustavo Bezerra for many conversations on viral proteinase mechanisms, and Jan Carette, Peter Sarnow, Gavin Sherlock and Priscilla Yang for insightful comments on the manuscript.

We are grateful for funding from NIAID (U19.AI171399, John Chodera, PI) and from Chem-H and Innovation grants from Stanford School of Medicine.

## MATERIALS AND METHODS

### Chemicals and Reagents

Telaprevir and Rupintrivir were purchased from ChemSpace. zVAMfmk was custom synthesized by Cortex Organics. Guanidine Hydrochloride was obtained from Sigma Aldrich and suspended in water. All other compounds were suspended in DMSO. Rabbit monoclonal anti-eIF4G antibody was purchased from Cell Signaling Technologies.

### Protease Constructs

Protease construct sequences from polio were derived from poliovirus type 1 Mahoney, EV71 sequences from an EV-A71 4643, and EV-D68 US MO/14/18947. Relevant sequences were amplified via PCR with the addition of BamHI or EcoRI and XhoI cleavage sites. Protease constructs were cloned into pT7CFE1-Chis (ThermoFisher) via T4 ligation (NEB). All mutations were made using QuikChange Lightning Mutagenesis kit (Agilent). Complete list of constructs and primers can be found in Table S2. EV-D68 US/MO/14-18927 WT infectious clone and EV-A71 4643 WT, 2A C110A, and 3C C147A infectious clones were a gift of Jan Carette.

### Translation Extracts

Translation reactions were performed using the TNT Quick Coupled Transcription/Translation kit (Promega). 10-50 ml reactions were assembled according to kit directions and labelled using EasyTag L-[35S]-Methionine (Perkin-Elmer). For pulse-chase reactions, after 20-30 minutes L-methionine was added to a final concentration of 1 mM and the temperature was raised to 37 °C to inhibit further translation. For dilution experiments, reactions were diluted with appropriate volumes of translation extract. At the indicated timepoint, aliquots were immediately mixed 1:1 with 2X Laemmli sample buffer. Samples were heated to 60 °C for 5 minutes and separated by SDS/PAGE. Gels were prepared and imaged as in (20) and quantified using ImageStudio (LI-COR). For experiments evaluating compound activity, either DMSO or the compound of interest was added at the beginning of the reaction. DMSO comprised 1% of the final reaction volume unless otherwise noted. Inhibition was analyzed after 30 minutes in the chase phase.

### Western Blots

For protease activity by in vitro translated protease, reactions were set up as above, with L-methionine substituted for EasyTag methionine. After 90 minutes, reactions were mixed 1:1 with 2X Laemmli sample buffer. To produce infected cell lysates, cells were infected at an MOI of 10 with poliovirus Type I Mahoney, incubated for 5 hours, and lysed by addition of 1X Laemmli buffer directly to cell monolayers. Both lysates and in vitro translation products were heated to 60 C for 5 min and separated by SDS/PAGE. Gels were transferred to PVDF membrane using BioRad TransBlot Turbo system, and the membrane was dried overnight. After rehydration, membranes were blocked for 1 hour with Intercept Blocking buffer TBS (LI-COR), and incubated overnight at 4 C with rabbit anti-eIF4G diluted 1:1000 in Intercept Blocking buffer. Membranes were then incubated for 1 hour with IRDye 800RD goat anti-Rabbit antibody diluted 1:10000 in Intercept buffer and imaged on a LI-COR Odyssey Fc Imager.

### Viral RNA cotransfection

HeLa or RD cells were electroporated as in (55) with the following modifications. Cells were resuspended to a concentration of 8×10^6^ cells/ml in DPBS, and 1 ml was added to the electroporation cuvette. 50 ug of RNA total was transfected into cells, at a ratio of 10:1 mutant or tRNA to WT viral RNA. Cells were electroporated with a Biorad GenePulser Xcell using an exponential decay protocol (300V, 500µF, ∞Ω). Cells were recovered in DMEM and aliquoted as necessary. Poliovirus cotransfections were performed in HeLa cells, and EV-A71 and EV-D68 cotransfections were performed in RD cells.

### Plaque Assay

After cotransfection, cells were aliquoted into 4 wells for technical replicates. After 12 hours (poliovirus), 16 hours (EV-A71), or 24 hours (EV-D68), cells were scraped into media and pelleted for 1 minute at 1000g. Cells were then resuspended in PBS++ and freeze-thawed 3 times. Viral titers were analyzed via plaque assay as described previously (56), using an avicel overlay. Poliovirus and EV-A71 plaque assays were grown for 2 days at 37 °C, and EV-D68 plaque assays were grown for 5 days at 34 °C.

### Luciferase Assay

After cotransfection, cells were aliquoted into 8 wells. At the indicated timepoint, cells were scraped into media and pelleted for 1 minute at 1000g. After removal of supernatant cells were resuspended in 100 ul of freshly prepared 1X Renilla Luciferase Assay Lysis buffer (Promega). Samples were analyzed with the Renilla Luciferase Assay System (Promega) according to kit directions on a Promega GloMax® Microplate reader.

### Statistical Analysis

All experiments n=3 unless otherwise noted. Quantifications were analyzed by two-way ANOVA with Dunnett’s multiple comparisons test unless otherwise noted. For *trans* cleavage assays, comparisons are to the wild-type self-processing. For dilution sensitivity assays, comparisons are to the undiluted reaction. *p<0.05; **p<0.01; ***p<0.001; ****p<0.0001. IC_50_ calculations were performed by fitting the data to equation 1:

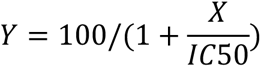

where X is the concentration of drug and Y is the percent cleavage. 100% cleavage is constrained to the amount of cleavage observed in the DMSO condition and 0% cleavage is defined as uncleaved precursor.

## SUPPLEMENTAL INFORMATION

**Table S1.**
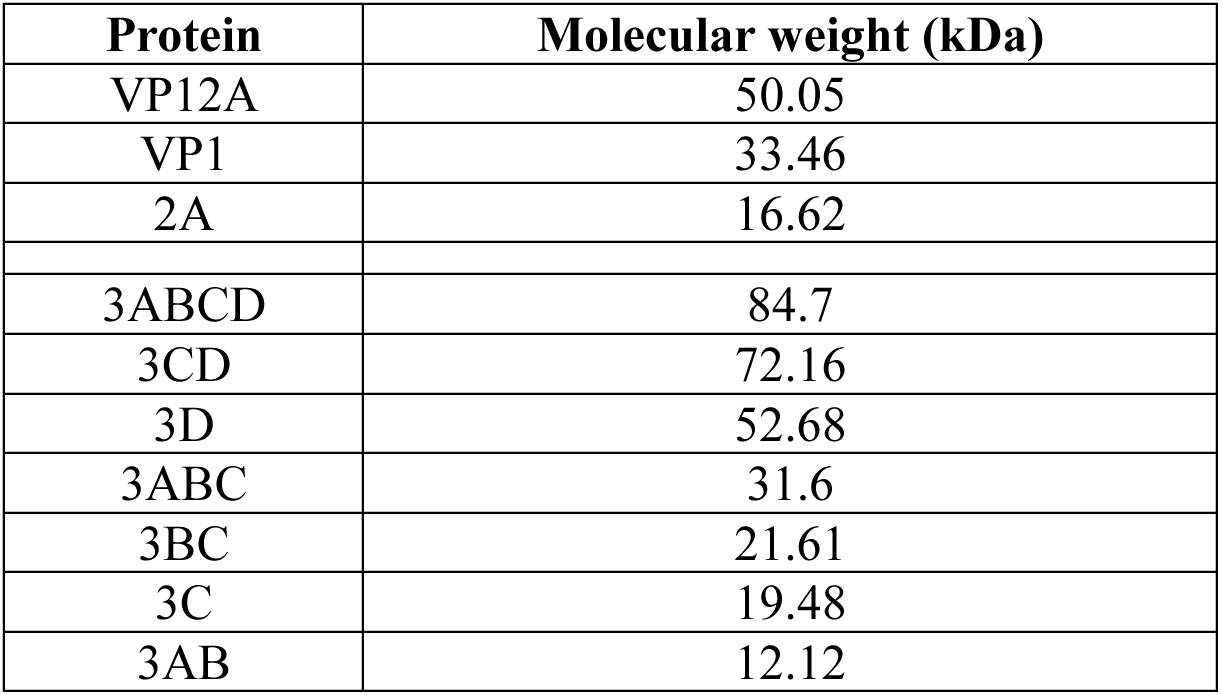
Molecular weights of proteins identified by SDS-PAGE.

**Figure S1.**
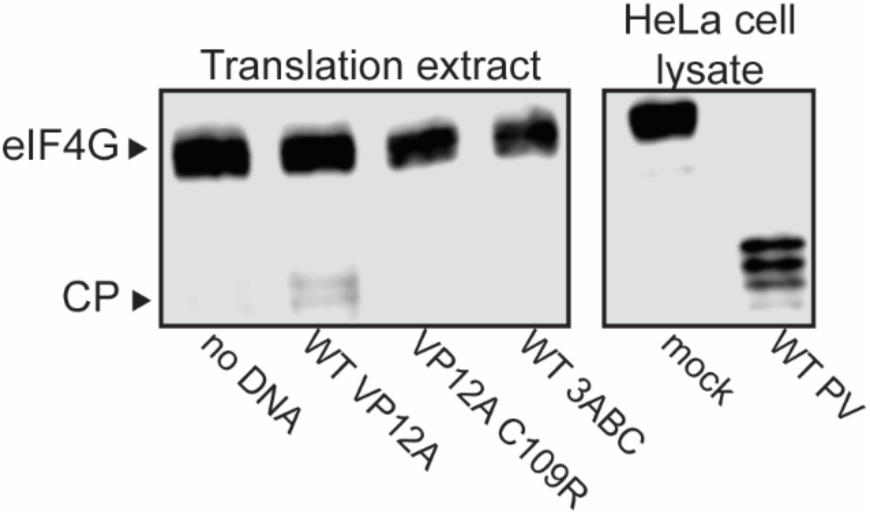
Poliovirus 2A made in translation extracts can cleave rabbit eIF4G. Translation extracts were programmed with the indicated DNAs. For infections, HeLa cells were either mock infected, or infected with poliovirus type 1 Mahoney. eIF4G abundance was visualized by immunoblotting, using an antibody that recognizes both human and rabbit eIF4G.

**Figure S2.**
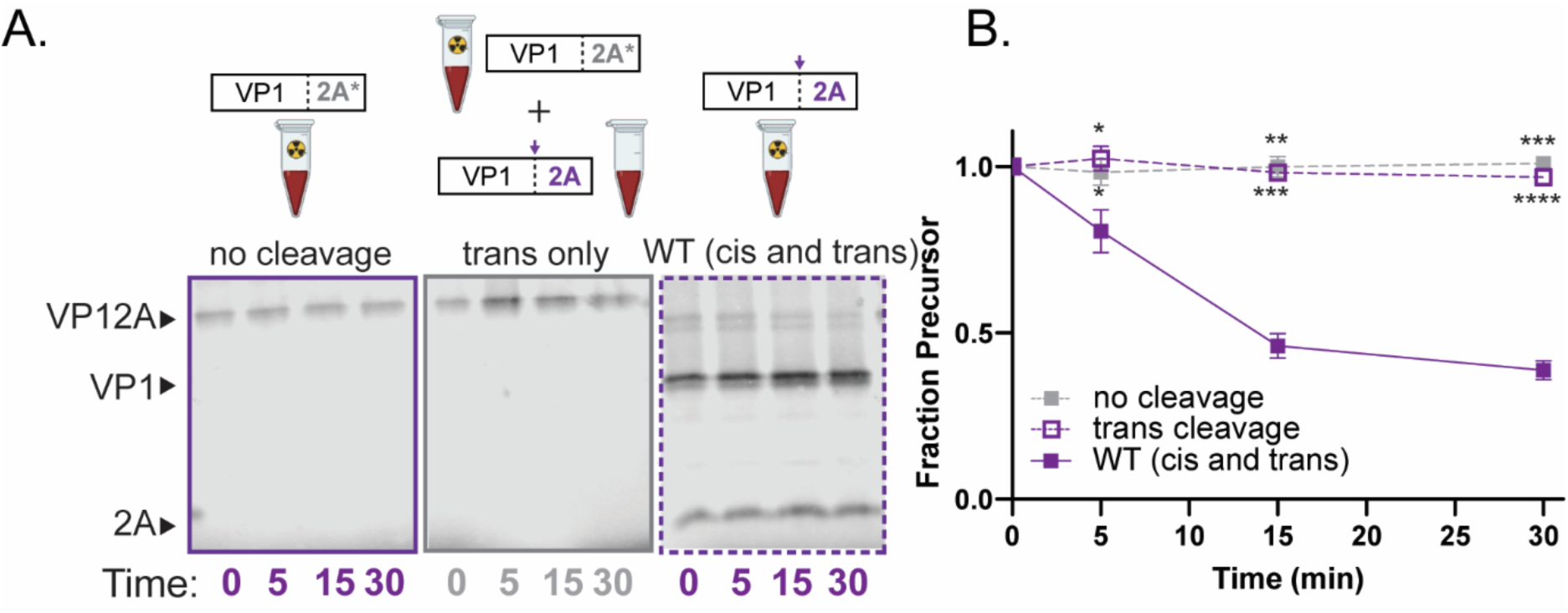
EV71 VP12A processing occurs in *cis*. Representative gels (A) and quantification (B) for EV-A71 VP12A *trans* cleavage assay. 2A* represents the C110A active site mutation. No cleavage: EV-A71 2A C110A with unlabeled 3ABC (negative control). *Trans* only: EV-A71 2A C110A with unlabeled catalytically active VP12A; WT (*cis* and *trans*): EV-A71 catalytically active VP12A.

**Figure S3.**
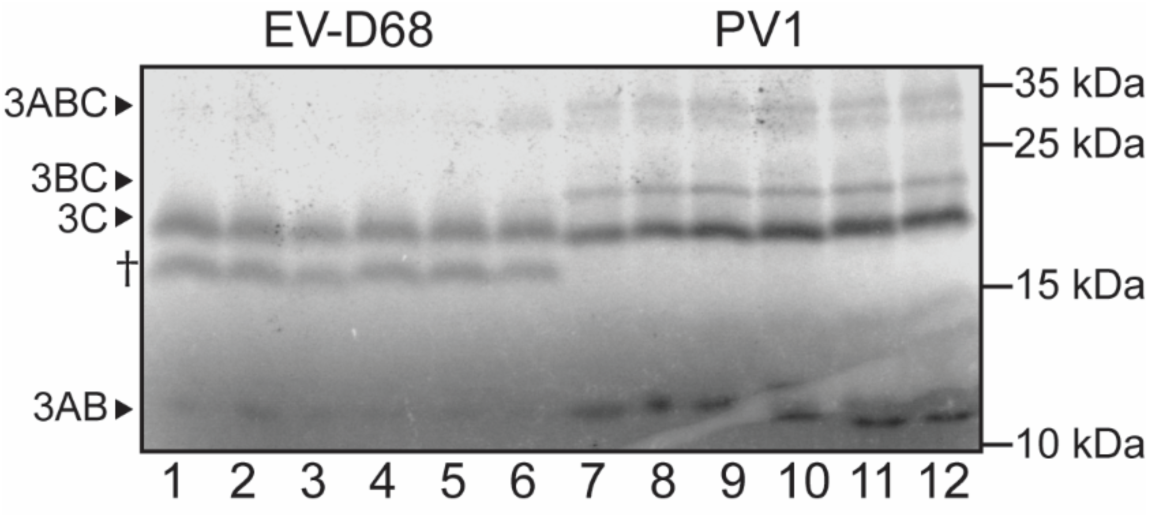
EV-D68 3ABC and 3ABCD cleavage produce an unidentified band. Lanes 1-6, translation extracts programmed with EV-D68 3ABC, pulsed for 30 minutes, and chased for 30 minutes. Lanes 7-12, translation extracts programmed with poliovirus 3ABC, pulsed for 30 minutes, and chased for 30 minutes. Unidentified band indicated with †. Band is approximately 16 kDa.

**Figure S4.**
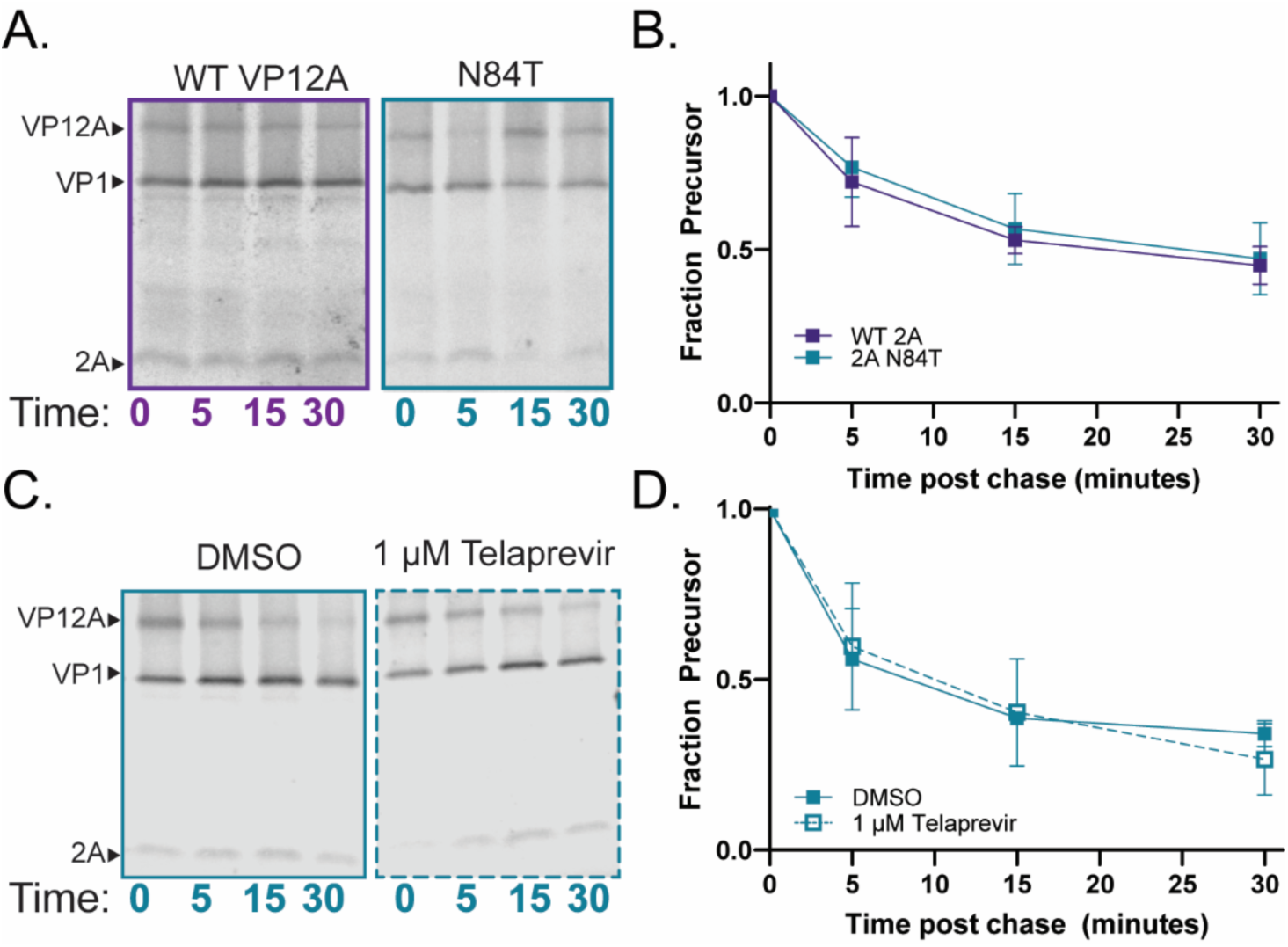
N84T mutant self-processing is not inhibited by telaprevir. (A and B) Representative gels (A) and quantification (B) of EV-D68 VP12A N84T mutant compared to wildtype processing. (C and D) Representative gels (C) and quantification (D) of N84T pulse chase in the presence of telaprevir. N84T mutation was previously characterized as conferring telaprevir resistance.(38)

**Figure S5.**
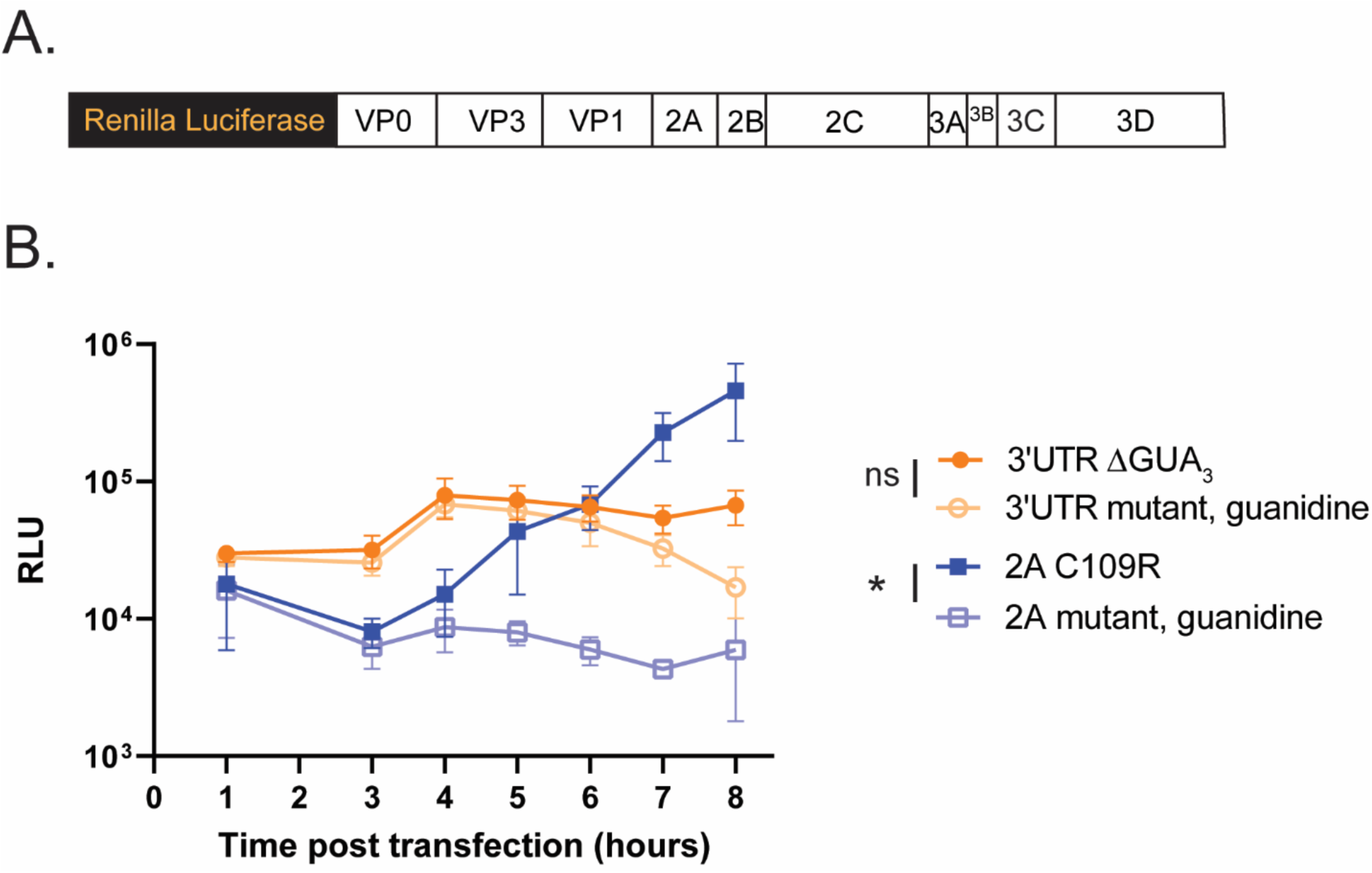
Genomes with mutations in the 2A protease active site can undergo RNA replication. A) Renilla luciferase virus construct. Luciferase gene connected to the N terminus of P1 via a 3C cleavage site. B) Cells were cotransfected with WT viral RNA and the indicated mutant luciferase construct. Luciferase signal is proportional to the amount of translation and RNA replication of the mutant viral construct.

**Table S2.**
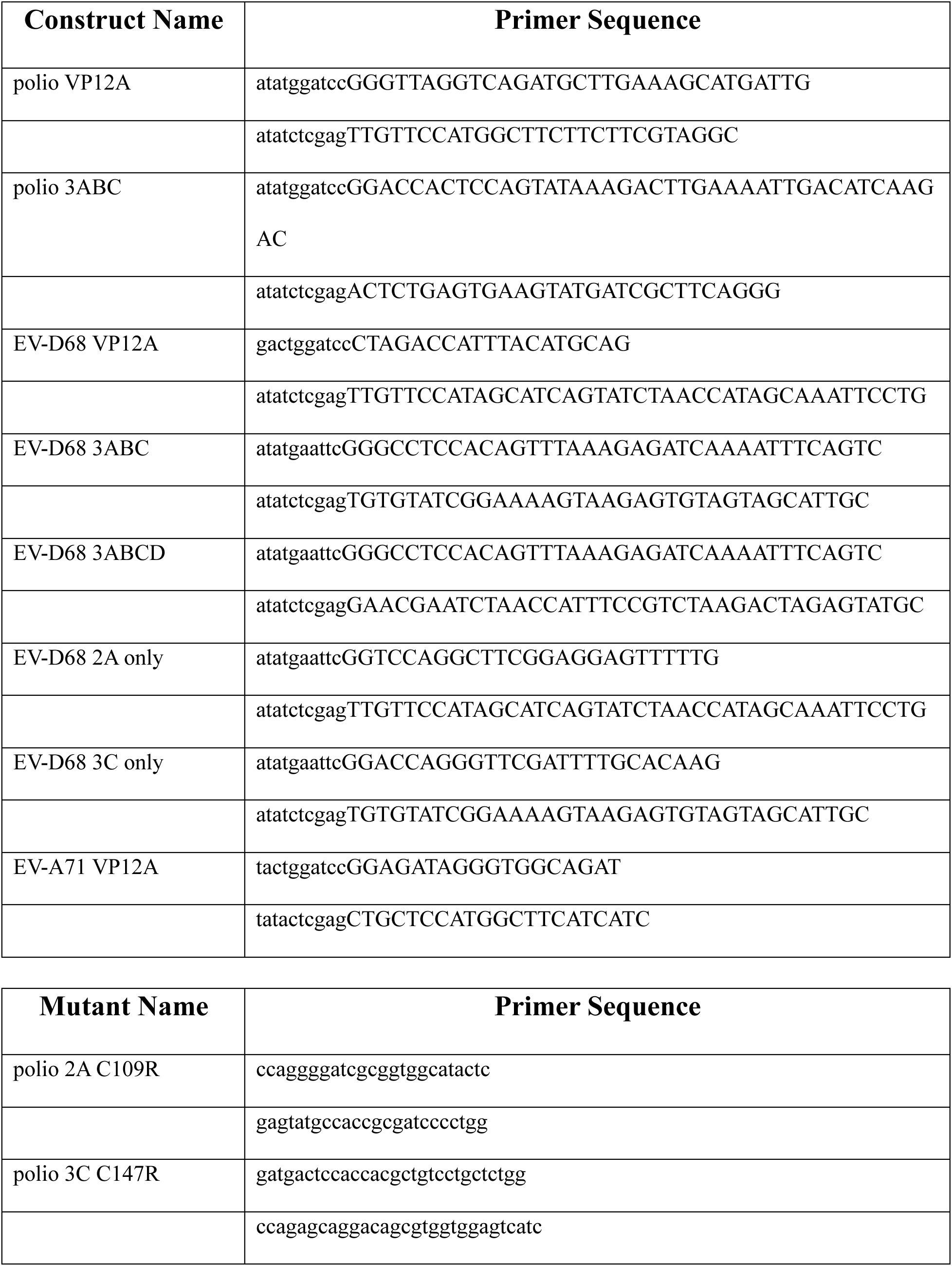
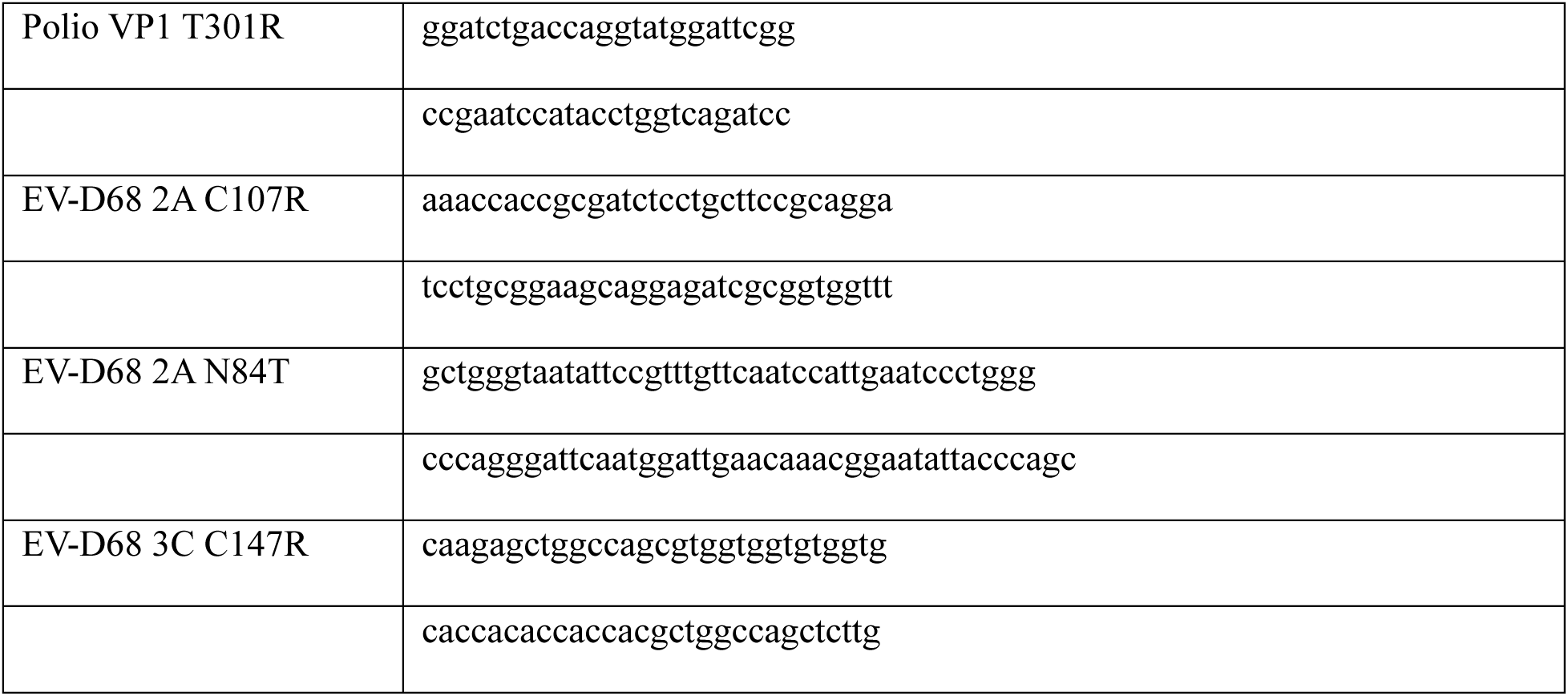
Primers used to make minimal protease constructs (top) and mutants (bottom). All primers ordered from IDT.

